# The genome design suite: enabling massive in-silico experiments to design genomes

**DOI:** 10.1101/681270

**Authors:** Oliver Chalkley, Oliver Purcell, Claire Grierson, Lucia Marucci

## Abstract

**Motivation:** Computational biology is a rapidly developing field, and *in-silico* methods are being developed to aid the design of genomes to create cells with optimised phenotypes. Two barriers to progress are that *in-silico* methods are often only developed on a particular implementation of a specific model (e.g. COBRA metabolic models) and models with longer simulation time inhibit the large-scale *in-silico* experiments required to search the vast solution space of genome combinations.

**Results:** Here we present the genome design suite (PyGDS) which is a suite of Python tools to aid the development of *in-silico* genome design methods. PyGDS provides a framework with which to implement phenotype optimisation algorithms on computational models across computer clusters. The framework is abstract allowing it to be adapted to utilise different computer clusters, optimisation algorithms, or design goals. It implements an abstract multi-generation algorithm structure allowing algorithms to avoid maximum simulation times on clusters and enabling iterative learning in the algorithm. The initial case study will be genome reduction algorithms on a *whole-cell* model of *Mycoplasma genitalium* for a PBS/Torque cluster and a Slurm cluster.

**Availability:** The genome design suite is written in Python for Linux operating systems and is available from GitHub on a GPL open-source licence.

**Contact:**, and.

## 1 Introduction

Mathematical models of cellular processes have been widely developed over the past 50 years to better understand and predict cellular behaviours; more recently, thanks to advances in Synthetic Biology, computational models can further aid in the rational design of cellular behaviours and, possibly, of entire genomes [1–21]. A drive to push biological models from specialising in specific processes to incorporate a more systems-level view of cells has led to transcription and translation being integrated into genome-scale metabolic models [22–26] and even the first whole-cell model [27]. The first whole-cell model is a first attempt to account for all annotated gene functions and molecular interactions of a cell in a single model and been validated on a broad range of data [27–29]. During this progression simulation times have increased from around one second in the case of genome-scale metabolic models to 5-35 hours in the case of the whole-cell model of *Mycoplasma genitalium* (*M. genitalium*). This means that running a genome design method on a new model requires one to recreate the method on the new model which can be time-consuming if not coded for general use or the new model is significantly different to use. Additionally, machine learning algorithms, like genetic algorithms used to optimse a phenotype by knocking-out combinations of genes [19, 30, 31], that run in reasonable amounts of time for simpler/faster models may take a prohibitively long time on more complex models - especially when taking into account that computer clusters often have a maximum simulation time. Furthermore these algorithms often require large numbers of simulations creating the need for relation database management systems and in cases where a simulation produces large amounts of data (e.g. the first whole-cell model [27]), distributed data storage solutions. Massive *in-silico* experiments with models as complex as the first whole-cell model become bigger than simply submitting a job to a cluster and so workflow becomes prohibitively fragmented and time consuming.

The area of workflow management systems (WMSs) arose in the 1970s as a way to automate workflows over distributed computing resources and now cover a huge range of different tasks for different types of organisations [32]. Scientific workflows [33, 34] have become more and more important as researchers gain access to computer clusters, cloud resources, remote databases, and even local databases that are prohibited access to the compute nodes of a cluster. The goals of the scientific workflow community were to create WMSs that hide all the complexity of distributed computing, allowing scientists to focus on the science and not the underlying infrastructure. Now scientific WMSs boast and impressive array of advanced features like easy to use graphical user interfaces, in-simulation monitoring and anomaly detection. However, this is very hard to generalise since even similar organisations or clusters will have their own bespoke infrastructure and corresponding security and process requirements which leads to very complicated set-up processes. The set-up process creates significant barriers of use to a user that does not have the full commitment of the organisation’s technology team to utilise a WMS. Additionally, there are many scientific workflow managers which are aimed at certain types of researchers by opening up the complexity of a certain aspect of the workflow (e.g. big data pipelines) making it very hard for a user to know which one to use. The variety of options combined with the high set-up cost create large barriers to the adoption of traditional scientific workflow managers.

Scripts of computer code often result in programs that do very specific tasks, and code must be rewritten if the user would like to do a slightly different task. Object-oriented programming is a programming paradigm based around grouping code into modules or objects [35, 36]. This can be done so that an object is a conceptual entity which can help humans more easily comprehend the code. Additionally, more extensive programs can be built from the fundamental modules. Object oriented-programming is often used to write complex generalisable code and was used to create the first ever whole-cell model [27].

## 2 PyGDS framework

Here we introduce PyGDS, a suite of computational tools which allows massive *in-silico* genome experiments to be performed on computer clusters thus overcoming time limits and workflow problems caused by complex models like the first whole-cell model [27]. This approach differs from traditional WMSs in that it does not attempt to be a fully functional system that hides lots of complexity to the user. In this case we provide a framework that enables communication between computers and the implementation of iterative algorithms that use the computers to implement simulations and perform data processing. The framework is written in Python, an easy to use and well known language, and the code is restricted to the most basic and fundamental functions like writing files and transferring data. This means that the user will have to write code to use the framework but it will be easy and transparent to understand the existing code. In essence, providing a simplified framework increases the amount of coding a user may have to do but allows them complete understanding of the code and thus autonomy from the help of third parties. PyGDS utilises the object-oriented paradigm to make it easier to modify the code for different computer clusters and algorithms. PyGDS is designed to run on a standard Linux PC that performs large-scale *in-silico* experiments by running batches of simulations on remote computer clusters; the standard computer will be referred to as the *hub* (an example set-up can be seen in figure 3 of the supplementary information).

To enable PyGDS to be adaptable to different design goals, models, and computer clusters the code is modularised in an object-oriented programming fashion. Three broad modules were picked: (i) algorithm, which provides a framework for implementing different algorithms on a computer model; (ii) computer communication, which provides a framework for communicating with a computer cluster, and (iii) job manager, which utilises the framework provided by the previous two modules to implement the algorithm onto computer clusters. For more information about how the modules integrate see figure 1 in the supplementary information.

**Figure 1:**
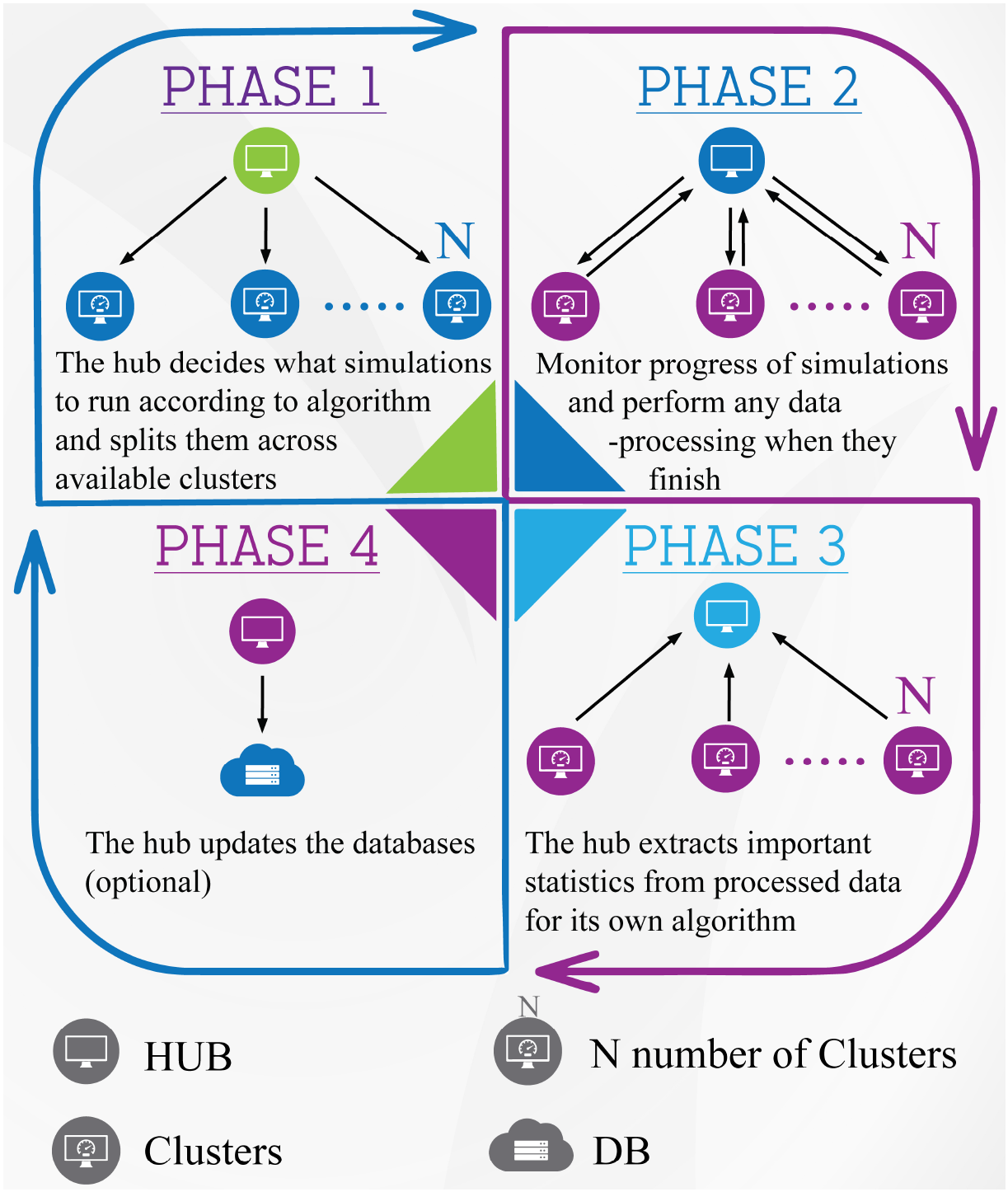
A diagram showing how PyGDS enables algorithms to be implemented on models across computer clusters.

The algorithm module utilises the idea of a multi-generation algorithm (MGA) to provide a framework for algorithms to be built on. A MGA describes a class of algorithms that iteratively runs *in-silico* experiments; each iteration is called a generation. The MGA has three benefits: (1) It is abstract, allowing many different algorithms to be implemented on a single framework (i.e. any algorithm that can be implemented on a given model in 1 or more generations); (2) provided that one generation can be simulated with the resources provided by the computer cluster(s), then the size and lifetime of the cluster is the only restriction on resource availability; (3) the iterative nature of a MGA allows results from previous generations to affect the decisions of future generations providing a mechanism for memory and learning in the algorithms. The computer communication module provides a framework to store all the data required to communicate with a remote Linux computer. The job submission module enables the algorithm module to run large batches of simulations on remote computer clusters by communicating with them through the computer communication module. Figure 1 shows how the modules integrate to perform large-scale *in-silico* experiments on a remote computer cluster(s).

For our case study we implement a genetic algorithm that reduces the genome of the whole-cell model of *M. genitalium* [27] using a cluster with a PBS/Torque job manager (i.e. BlueCrystalIII), and cluster with a Slurm job manager (i.e. BlueGem) - see figure 3 of the supplementary information. Both job managers are commonly used for computer clusters and so these examples can be used as templates for other clusters with these job managers, otherwise they still act as an example of how to write a `connection` sub-class - for more information on this see the supplementary information. For an example of implementing a different algorithm using PyGDS see the guess, add, mate algorithm (GAMA) in [37] which is implemented as three separate algorithms using the PyGDS framework where the results of a previous stage are fed into the next next stage - combined these three stages act like a modified genetic algorithm that is designed to converge on a minimal genome faster.

In order to develop genome design algorithms, subclasses from each of the three modules need to be created allowing the framework to know what model to use on which cluster(s) and the details of the algorithm. Projects that have already been developed can be adjusted to run on a different cluster or algorithm by creating an appropriate subclass. For more information on how this was done please see the supplementary information.

## 3 Conclusion

PyGDS provides a framework to implement multi-generation algorithms on (potentially multiple) computer clusters and distributed storage systems. The case study shows how to implement a genetic algorithm and run it on different clusters. Rees and Chalkley et al. [37] also show how to implement a different algorithm. Whilst it has not been shown yet, the modularity of the code may make it possible to use different models and even different design goals. The genome design suite helps to overcome limitations of maximum simulation times on clusters, restriction to a single computer cluster, and takes a step towards providing a framework to develop genome design tools that are not restricted to specific models or clusters. Whilst being more general in principle, the purpose of PyGDS is to allow the optimisation of phenotypes of *in-silico* organisms. In the future it may be possible to test algorithms against different models/organisms or perform massive *in-silico* experiments on simpler models to simulate the interaction of large communities of cells. The fact that different models and design goals are possible open up uses outside of genome design like hyper-parameter optimisation or parameter fitting and could even be used on non-biological models. However, the further a user moves from the case study the more code they have to write. Whilst it is modularised well for variable computer clusters and algorithms it is not so good for different models and design goals - improvement of this is expected to the center of the next version update. If a user is able to confine an entire project to one computer or a single cloud services provider then there is no need for a WMS but as soon as workflows become distributed then a manager becomes useful. Additionally, whilst cloud services provide almost unlimited computing facilities to a user there is a fee for data storage and CPU/GPU usage and so if that user has access to cheaper resources (i.e. an in-house server, cluster, or data storage facilities) then a WMS can be used to maximise the use of the cheaper resources and minimise the use of more expensive resources.

## Supporting information

The whole supplementary information document.

## Acknowledgements

I would like to thank Dr Jonathan Karr and Prof. Markus Covert for all advice given with regards to the whole-cell model of *M. genitalium*. I would also like to thank Joshua Rees and Sophie Landon of the genome design group at the University of Bistol for all advice and testing done throughout the development of PyGDS. I would like to thank the ACRC at the University of Bristol for access to BlueCrystal, BlueGem, and other high-performance computing facilities as well invaluable advice from Steve Roome, Simon Burbidge, Matthew Williams, and Dr Christopher Woods.

## References

[1] Kiran Patil, Isabel Rocha, Jochen Förster, and Jens Nielsen. Evolutionary programming as a platform for in silico metabolic engineering. BMC Bioinformatics, 6(1):308, dec 2005.

[2] Yohei Shinfuku, Natee Sorpitiporn, Masahiro Sono, Chikara Furusawa, Takashi Hirasawa, and Hiroshi Shimizu. Development and experimental verification of a genome-scale metabolic model for Corynebacterium glutamicum. Microbial cell factories, 8:43, aug 2009.

[3] J S Edwards and B O Palsson. The Escherichia coli MG1655 in silico metabolic genotype: its definition, characteristics, and capabilities. Proceedings of the National Academy of Sciences of the United States of America, 97(10):5528–33, may 2000.

[4] Kiran Raosaheb Patil, Mats Åkesson, and Jens Nielsen. Use of genome-scale microbial models for metabolic engineering. Current Opinion in Biotechnology, 15(1):64–69, feb 2004.

[5] Bashir Sajo Mienda. Genome-scale metabolic models as platforms for strain design and biological discovery, 2017.

[6] Aarash Bordbar, Jonathan M Monk, Zachary A King, and Bernhard O Palsson. Constraint-based models predict metabolic and associated cellular functions. Nature reviews. Genetics, 15(2):107–20, feb 2014.

[7] Adam M Feist, Daniel C Zielinski, Jeffrey D Orth, Jan Schellenberger, Markus J Herrgard, and Bernhard Ø Palsson. Model-driven evaluation of the production potential for growth-coupled products of Escherichia coli. Metabolic engineering, 12(3):173–86, may 2010.

[8] Axel von Kamp and Steffen Klamt. Enumeration of smallest intervention strategies in genome-scale metabolic networks. PLoS computational biology, 10(1):e1003378, jan 2014.

[9] Stefan Schuster, Thomas Dandekar, and David A. Fell. Detection of elementary flux modes in biochemical networks: A promising tool for pathway analysis and metabolic engineering. Trends in Biotechnology, 17(2):53–60, 1999.

[10] Joonhoon Kim and Jennifer L Reed. RELATCH: relative optimality in metabolic networks explains robust metabolic and regulatory responses to perturbations. Genome biology, 13(9):R78, jan 2012.

[11] Chiam Yu Ng, Moo-young Jung, Jinwon Lee, and Min-Kyu Oh. Production of 2,3-butanediol in Saccharomyces cerevisiae by in silico aided metabolic engineering. Microbial Cell Factories, 11(1):68, 2012.

[12] Harry Yim, Robert Haselbeck, Wei Niu, Catherine Pujol-Baxley, Anthony Burgard, Jeff Boldt, Julia Khandurina, John D Trawick, Robin E Osterhout, Rosary Stephen, Jazell Estadilla, Sy Teisan, H Brett Schreyer, Stefan Andrae, Tae Hoon Yang, Sang Yup Lee, Mark J Burk, and Stephen Van Dien. Metabolic engineering of Escherichia coli for direct production of 1,4-butanediol. Nature chemical biology, 7(7):445–52, jul 2011.

[13] Mounir Izallalen, Radhakrishnan Mahadevan, Anthony Burgard, Bradley Postier, Raymond Didonato, Jun Sun, Christopher H Schilling, and Derek R Lovley. Geobacter sulfurreducens strain engineered for increased rates of respiration. Metabolic engineering, 10(5):267–75, sep 2008.

[14] Jin Hwan Park, Kwang Ho Lee, Tae Yong Kim, and Sang Yup Lee. Metabolic engineering of Escherichia coli for the production of L-valine based on transcriptome analysis and in silico gene knockout simulation. Proceedings of the National Academy of Sciences of the United States of America, 104(19):7797—802, may 2007.

[15] Kento Tokuyama, Satoshi Ohno, Katsunori Yoshikawa, Takashi Hirasawa, Shotaro Tanaka, Chikara Furusawa, and Hiroshi Shimizu. Increased 3-hydroxypropionic acid production from glycerol, by modification of central metabolism in Escherichia coli. Microbial cell factories, 13:64, jan 2014.

[16] Sang Jun Lee, Dong-Yup Lee, Tae Yong Kim, Byung Hun Kim, Jinwon Lee, and Sang Yup Lee. Metabolic engineering of Escherichia coli for enhanced production of succinic acid, based on genome comparison and in silico gene knockout simulation. Applied and environmental microbiology, 71(12):7880—7, dec 2005.

[17] Hal Alper, Yong-Su Jin, J F Moxley, and G Stephanopoulos. Identifying gene targets for the metabolic engineering of lycopene biosynthesis in Escherichia coli. Metabolic engineering, 7(3):155—64, may 2005.

[18] Ana Rita Brochado, Claudia Matos, Birger L Møller, Jørgen Hansen, Uffe H Mortensen, and Kiran Raosaheb Patil. Improved vanillin production in baker’s yeast through in silico design. Microbial cell factories, 9(1):84, jan 2010.

[19] Mohammad A Asadollahi, Jérôme Maury, Kiran Raosaheb Patil, Michel Schalk, Anthony Clark, and Jens Nielsen. Enhancing sesquiterpene production in Saccharomyces cerevisiae through in silico driven metabolic engineering. Metabolic engineering, 11(6):328–34, nov 2009.

[20] Douglas McCloskey, Bernhard Ø Palsson, and Adam M Feist. Basic and applied uses of genome-scale metabolic network reconstructions of Escherichia coli. Molecular systems biology, 9:661, jan 2013.

[21] Edward J. O’Brien, Jonathan M. Monk, and Bernhard O. Palsson. Using Genome-scale Models to Predict Biological Capabilities. Cell, 161(5):971–987, may 2015.

[22] M W Covert, C H Schilling, and B Palsson. Regulation of gene expression in flux balance models of metabolism. Journal of theoretical biology, 213(1):73–88, nov 2001.

[23] Timothy E Allen and Bernhard Ø Palsson. Sequence-based analysis of metabolic demands for protein synthesis in prokaryotes. Journal of theoretical biology, 220(1):1–18, jan 2003.

[24] Edward J O’Brien, Joshua A Lerman, Roger L Chang, Daniel R Hyduke, and Bernhard Ø Palsson. Genome-scale models of metabolism and gene expression extend and refine growth phenotype prediction. Molecular systems biology, 9:693, jan 2013.

[25] Ines Thiele, Ronan M T Fleming, Richard Que, Aarash Bordbar, Dinh Diep, and Bernhard O Palsson. Multiscale modeling of metabolism and macromolecular synthesis in E. coli and its application to the evolution of codon usage. PloS one, 7(9):e45635, jan 2012.

[26] Colton J. Lloyd, Ali Ebrahim, Laurence Yang, Zachary A. King, Edward Catoiu, Edward J. O’Brien, Joanne K. Liu, and Bernhard O. Palsson. COBRAme: A computational framework for genome-scale models of metabolism and gene expression. PLoS Computational Biology, 14(7), 2018.

[27] Jonathan R Karr, Jayodita C Sanghvi, Derek N Macklin, Miriam V Gutschow, Jared M Jacobs, Benjamin Bolival, Nacyra Assad-Garcia, John I Glass, and Markus W Covert. A whole-cell computational model predicts phenotype from genotype. Cell, 150(2):389–401, jul 2012.

[28] Jayodita C Sanghvi, Sergi Regot, Silvia Carrasco, Jonathan R Karr, Miriam V Gutschow, Benjamin Bolival, and Markus W Covert. Accelerated discovery via a whole-cell model. Nature methods, 10(12):1192–5, dec 2013.

[29] Oliver Purcell, Bonny Jain, Jonathan R Karr, Markus W Covert, and Timothy K Lu. Towards a whole-cell modeling approach for synthetic biology. Chaos (Woodbury, N.Y.), 23(2):025112, jun 2013.

[30] Kiran Raosaheb Patil, Isabel Rocha, Jochen Förster, and Jens Nielsen. Evolutionary programming as a platform for in silico metabolic engineering. BMC bioinformatics, 6:308, jan 2005.

[31] Isabel Rocha, Paulo Maia, Pedro Evangelista, Paulo Vilaça, Simão Soares, José P Pinto, Jens Nielsen, Kiran R Patil, Eugenio C Ferreira, and Miguel Rocha. OptFlux: an open-source software platform for in silico metabolic engineering. BMC systems biology, 4(1):45, jan 2010.

[32] Edward A. Stohr and J. Leon Zhao. Workflow Automation: Overview and Research Issues. Information Systems Frontiers, 3(3):281–296, 2001.

[33] Bertram Ludäscher, Ilkay Altintas, Chad Berkley, Dan Higgins, Efrat Jaeger, Matthew Jones, Edward A. Lee, Jing Tao, and Yang Zhao. Scientific workflow management and the Kepler system. Concurrency and Computation: Practice and Experience, 18(10):1039–1065, aug 2006.

[34] Ewa Deelman, Tom Peterka, Ilkay Altintas, Christopher D Carothers, Kerstin Kleese van Dam, Kenneth Moreland, Manish Parashar, Lavanya Ramakrishnan, Michela Taufer, and Jeffrey Vetter. The future of scientific workflows. The International Journal of High Performance Computing Applications, 32(1):159–175, jan 2018.

[35] Ole-Johan Dahl and Kristen Nygaard. Class and Subclass Declarations. In Software Pioneers, pages 91–107. Springer Berlin Heidelberg, Berlin, Heidelberg, 2002.

[36] John C. Mitchell. Concepts in Programming Languages. Cambridge University Press, Cambridge, 2002.

[37] Joshua Rees, Oliver Chalkley, Sophie Landon, Oliver Purcell, Lucia Marucci, and Claire Grierson. Designing Minimal Genomes Using Whole-Cell Models. bioRxiv, page 344564, mar 2019.

