## Supplementary material for "The genome design suite: enabling massive in-silico experiments to design genomes": The whole supplementary information document.

### Supplementary materials for the genome design suite: enabling massive in-silico experiments to design genomes

[1]Engineering mathematics, University of Bristol, Bristol, BS8 3PF, UK.
[2]MIT Synthetic Biology Center, 500 Technology Square, NE47-257, Cambridge, MA 02139, USA.
[3]School of Biological Sciences, University of Bristol, Bristol, BS8 3PF, UK.
[4]Co-last author.

#### 1 PyGDS framework

PyGDS is a Python library created to provide a framework to enable massive *in-silico* experiments. All computers are assumed to have Linux operating systems. As outlined in the main text, three fundamental processes make up PyGDS:

1. Decide what simulations to run next in-order to optimise a specified function and learn from previous simulations if there is any;
2. organise simulations into batches and submit them to the computer cluster(s). Monitor all running jobs and when the jobs are finished, perform any necessary tasks like data processing and updating databases;
3. perform fundamental tasks on a remote computer like creating files, running code, and checking disk usage.

The fundamental processes are coded into the following modules:

- Process-1, or *algorithms*, is in the `multigeneration_algorithm.py` module.

*
†
‡

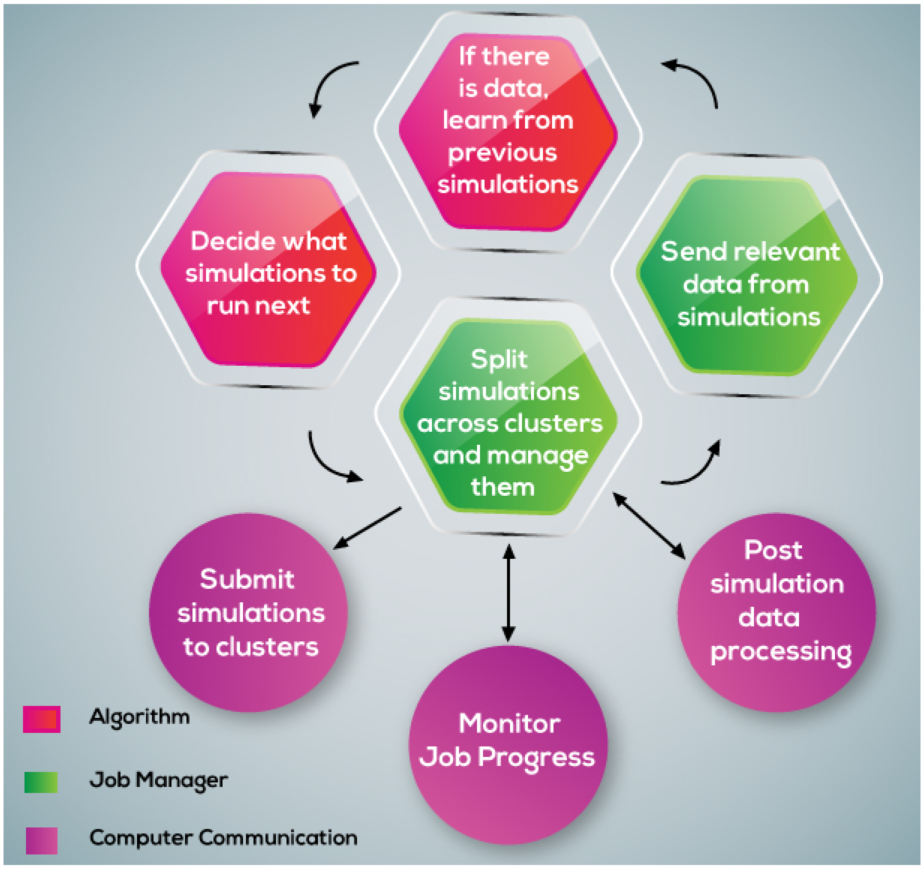


Figure 1: This shows how the three fundamental processes interact to design an *in-silico* genome. Process-1 is labelled as 'algorithm', process-2 is labelled as 'job manager', and process-3 is labelled as 'computer communication'.

- Process-2, or *job manager*, is in the `batch_jobs.py` module.
- Process-3, or *computer communication*, is in the `base_connections.py` and `connections.py` modules.

Supplementary Figure 1 shows how these processes interact to create an iterative process that learns over time. Furthermore, if coded correctly, these fundamental processes can act as an abstract template enabling versatility of algorithms, models and computer clusters.

#### 1.1 Computer communication

Generally, computer clusters can only be accessed by authorised users through a secure shell (SSH) remote connection. Authorised users are required to prove their identity by either password or encryption key. There is no universal login system for clusters and PyGDS will need repeated access, so it is required that a user has certain SSH settings. Coding passwords or storing them is insecure, and so a user's account must be accessible by encryption key (this is a standard method for logging into a computer cluster and is generally perceived to be more secure than using passwords). The user will then need to set up the following file ' /.ssh/config':

```
Host ssh_alias
        User user_name
        HostName address_to_remote_computer
        IdentityFile /home/user_name/.ssh/key_name
```

This setup means that in a Linux terminal it is possible to connect to the remote machine, without manually entering a password, using the following command `ssh ssh_alias` .

The computer communication code is kept in the `base_connection` module. The `Connection` class is the only class in the base_connection module and is the abstract class that defines the core structure of all connection classes. All the child classes that inherit from this class can be thought of as a portal to another computer. This document will provide two example subclasses BlueCrystalIII, and BlueGem which are both high-performance computer clusters based at the University of Bristol. The former uses a PBS/Torque job manager and the latter uses a Slurm job manager. These are both popular job managers for computer clusters and so can be used as a template for subclasses to other computer clusters.

Initialisation:
The `Connection` class is initialised with attributes:

- The username of the account on the remote computer.
- The SSH alias defined in ' /.ssh/config' file.

- The path to the encryption key required for user login.
- The forename of the user.
- The surname of the user.
- The user's email.

Instance methods:
The `createFile` method turns a list of text into a file on the local computer and sets the file permissions if specified.

The `rsyncFile` method transfers files either locally or onto the remote computer. This method uses the `rsync` function commonly found on Linux computers and can be installed with `apt-get install rsync` on Debian-based systems or `yum install rsync` on RPM-based systems.

The `convertKosAndNamesToFile` method is specific to gene knock-out experiments with the whole-cell model of *M. genitalium*[1]. It creates two files, one that contains all the sets of gene knock-outs and one that contains a name for each set. Both files are ordered in the same way, so line 1 of the names file gives the name of the knock-out set on line 1 of the gene knock-out sets file.

The `sendCommand` method takes a list of shell commands and runs them on the remote computer. It returns a dictionary with the `stdout` , `stderr` and `return_code` .

Static methods:
The `checkSuccess` method takes a function that needs to make a remote connection with a set of arguments and executes that function in a `while` loop until the return code signifies success. Having it loop normally can potentially overload the login server of the remote computer (like a denial of service attack), and so this waits a certain duration of time before trying again. It starts relatively frequent and slows down, starting with every three seconds and ending by checking once every 24 hours. Once the function returns a successful return code, `checkSuccess` returns the output with a successful return code. If the function does not return a successful return code within 7 days, then it exits with a return code of 13 but this can be easily changed to a user's preference by editing the source code or overloading the method in a sub-class of this.

Abstract methods:
`checkQueue` and `checkDiskUsage` are methods that will often be used but vary from computer to computer. When creating a child class for a new computer to connect to, it is advised that these methods are properly overloaded because they are made abstract so that additional software that uses the framework can assume that this functionality is available. If a method(s) is not available, then compilation or potentially dangerous runtime errors may occur. However,

anyone that does not wish to add this or the computer does not have the functionality can create overload the method with only the `pass` command so that it does nothing. The `checkQueue` method checks the queue on a remote cluster and the `checkDiskUsage` returns disk usage statistics of the remote computer.

#### 1.2 Job manager

There is no framework for the job manager section as the structure is provided by the `base_connection` and the `multigeneration_algorithm` modules. The job manager's tasks depend on the what data processing and storage solution the user desires and so these modules are only coded for specific implementations.

#### 1.3 Algorithms

The `MGA` class is an abstract class that acts as a template that all other algorithm classes should inherit from and can be found in the `multigeneration_algorithm` module. The class must be abstract enough that it can act as a template for as many algorithms as possible and can accommodate any single or multi-generation algorithm that is compatible with the model being used. Supplementary Figure 2 shows how the `MGA` class, and thus all algorithms, execute. One can see that all algorithms will be started by running the `run` method, which then initiates a loop that does not stop until a specified maximum generation number is reached. Each iteration of the loop represents a single generation and each generation is created and simulated using the `runSimulations` method.

<u>Initialisation</u>:
The class needs to be initialised with:

1 A Python dictionary where the values are cluster connection instances that are available to run simulations on (i.e. child classes of the `Connection` class from the `base_connections` module). The keys of the clusters are unique names that label each cluster connection.

2 A Python dictionary that defines when the algorithm should stop running. At present this only has the functionality to stop at a predetermined generation in the future, but additional options could be added.

3 The name given to this instance of the algorithm.

4 Once the class is initialised a class variable that remembers what generation the algorithm is on is created and set to `None` .

<u>Instance methods:</u>
The `checkStop` method checks to see if the generation counter (i.e. class variable 4) is less than the 'stop generation' number in the 'checkStop' dictionary (i.e. class variable 2). If the generation counter (4) is equal or more than the 'checkStop' dictionary (2) it returns `True` otherwise it returns `False` .

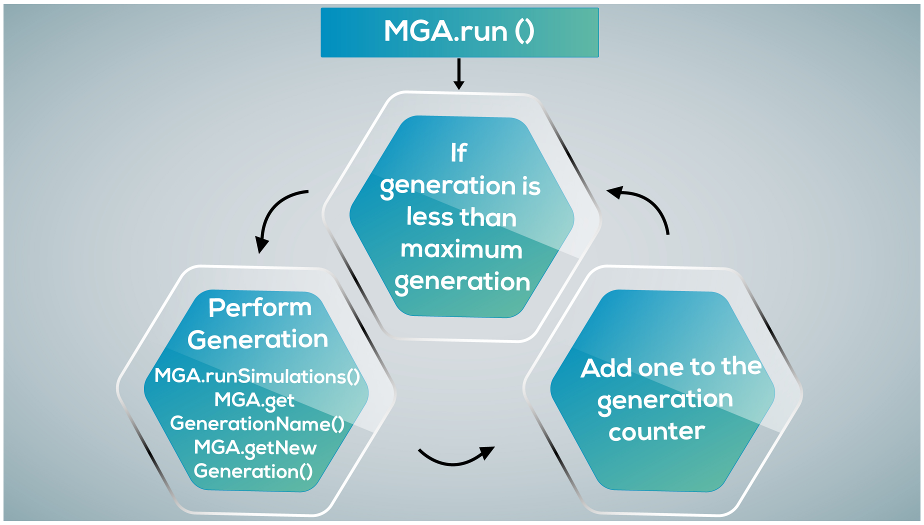


Figure 2: A schematic of the abstract class MGA. One can see that all algorithms will be started by using the `run()` method which initiates a loop that repeats until a maximum generation is reached. It is an abstract class, so needs a child class to be used and that child needs to define what the algorithm does through the `runSimultions()` , `getGenerationName()` , and `getGeneration()` methods.

The `run` method is a `while` loop that runs the next batch of simulations and increments the generation counter (i.e. class variable 4) by one until the generation counter is greater or equal to the maximum generation (i.e. `while checkStop()!= True` ).

<u>Abstract methods:</u>
The `runSimulations` method is called by the `run` method and implements one generation of an algorithm. Since this is the abstract class it does not specify any algorithm and leaves it for the child classes to define. Two other abstract methods are set as they will ned to be called by the `runSimulations` method. These are the `getGenerationName` and `getNewGeneration` methods.

The `getNewGeneration` method will need to create the genomes of the children that need to be simulated in the next generation but is an abstract method and so it is left for the child class to define.

The `getGenerationName` method will return a name for each generation that can be used to identify what belongs to each generation. This is an abstract method and so it is left for the child classe to define.

### 2 Implementation tutorial

In order to demonstrate the use of PyGDS we will use the case study from the main text. The case study wishes to use a genetic algorithm to reduce the genome of the whole-cell model of *M. genitalium.* In order to demonstrate using a PBS/Torque cluster and Slurm cluster the case study will use BlueCrystalIII and BlueGem to run the simulations, respectively. For an example of implementing a different algorithm see the supplementary information of the guess, add, and mate algorithm[2]. Here, we report firstly details of the resources at the University of Bristol, setting up access to the cluster, and details on running the whole-cell model are presented. Then, we include details of how to create both the `Connection` and the `MGA` sub-classes and writing the job management classes. Finally, the steps to starting the genetic algorithm are presented.

Genetic algorithms are a great general purpose, easy to implement machine learning algorithm used to optimise objectives and have been used to in a wide variety of tasks[3, 4]. A genetic algorithm attempts to, roughly, mimic evolution by natural selection in order to learn. The population of individuals is made up of parents and children where the parents are the fittest individuals that survived whatever *natural selection* is placed upon them. Here the natural selection is normally implemented by some kind of objective function that one wishes to optimise. A set of genes defines each parent and the fittest individuals *mate* to create children that are made up of a random combination of the genes of both parents, plus some random mutation.

In order for PyGDS to control clusters from the hub, the user must be able to log into the remote cluster without a password as described in section 1. Once this is achieved, the user must create subclasses of the `Connection` and `MGA` classes in order to utilise the framework. Then the job manager classes need writing so that the algorithm knows how to split simulations across the cluster(s), perform data processing, and update any databases - depending on the data requirements of the user. The rest of this section will give a general explanation of the process and will provide an example based on setting up a genetic algorithm to reduce the genome of the whole-cell model of *M. genitalium.* The requirements of this will be described in the next section followed by descriptions of sub-classing the `Connection` and `MGA` classes, writing the job management classes, and finally running the genetic algorithm on the computer cluster.

#### 2.1 Resources and requirements of the case study

The University of Bristol provides access to two high-performance computing clusters, BlueCrystalIII (BC3), BlueGem (BG), and the Flex1 scratch drive. The hub is an standard desktop computer (Intel(R) Core(TM) i3-3110M CPU @ 2.40GHz with 4GB RAM). Supplementary Figure 3 shows that the hub can access BC3 and BG and that the clusters have access to a shared scratch drive,

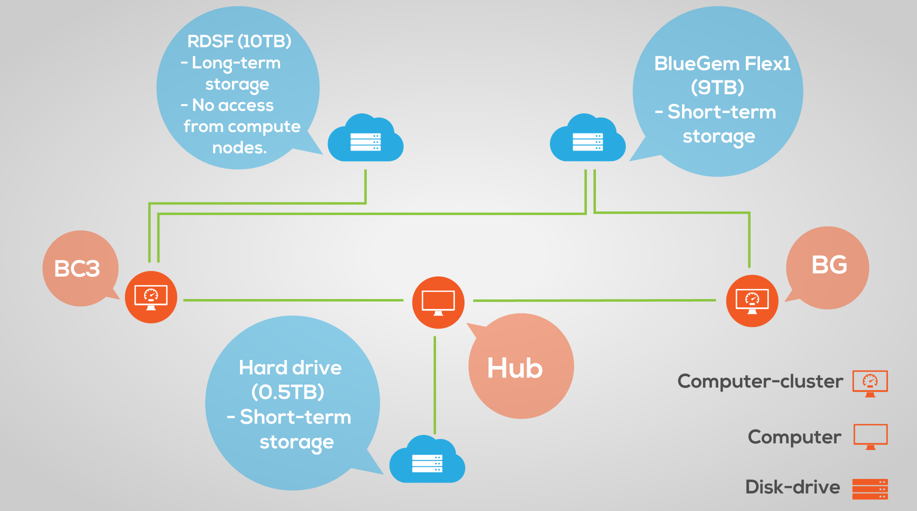


Figure 3: A diagram showing the resources at the University of Bristol and how they are connected.

flex1. The hub ran on the CentOS 6.6 distribution of Linux; BC3 runs Scientific Linux 6.4 (Carbon) and uses the PBS 4.2.4.1 job scheduler - for more information about BC3 see the official webpage at [5]; BG runs on Scientific Linux release 6.6 (Carbon) and uses the Slurm 14.03.0 - for more information see the official webpage at [6].

All simulation output data will be written to the shared scratch drive in two forms.

1. SQLite3 database called *ko.db* to store general details about each batch of simulations and the average growth rate and division time of each *in-silico* cell.

2. The raw simulation output data of a single simulation is split into hundreds of compressed Matlab files which are slow to access, and so it was decided to convert the data into Python Pandas Dataframes stored as Pickle files. In [2] GAMA ran 51,119 simulations which would produce $\sim$ 12.2TBs of data[1] and so some generally important time series are saved and the rest are deleted. These files are called 'basic_summary*.pkl' where * is some kind of identifier, and an example of a wild-type simulation is contained in SI.zip. The Pickle files contain the following time series.

    - Chromosome state: ploidy and segregation status.
    - FtsZ ring: variables related to the state of the division process (i.e.

[1]It was estimated that an average wild-type simulation produced $\sim$ 239MBs of data.

```
classdef koRunner < edu.stanford.covert.cell.sim.runners.
    SimulationRunner
        methods
                function this = koRunner(varargin)
                        this = this@edu.stanford.covert.cell.sim.
                            runners.SimulationRunner(varargin{:});
                end
        end

    % jobNumber can be used in order to change make the simulation
        jobNumber depended.
    % Change the jobnumber as shown in this example:
    % runSimulation(..,'runner','AdjustParameters','jobNumber
        ',1,..)
    properties
        jobNumber = 0;              % Preallocate jobNumber
        koList=0;
    end

        methods (Access = protected)
        function modifyNetworkParameters(this, sim)
            % set the seeed and display it
            timeNow = clock;
            seed = this.jobNumber * 10000 + timeNow(2)*24*31 +
                timeNow(3)*24 + timeNow(4);  %Override seed
            this.seed = seed

            % apply kos
            koList=this.koList
            sim.applyOptions('geneticKnockouts', koList)
        end
        end
end
```

Figure 4: The simulation runner sub-class used to perform gene knock-out experiments.

the number of edges and whether they are bent). For more information about how cell division is modelled, see [1].

- Geometry: the diameter of the cell and if the cell has completely separated into two cells.
- Mass: various metrics of mass (e.g. total, DNA, RNA, protein, metabolite, media, and water).
- Metabolic reactions: growth.
- Ribosome: states of individual ribosomes (e.g. active, stalled, or formed).
- RNA-polymerase: states of individual RNA-polymerase (e.g. active, free, specifically bound, and non-specifically bound).

```
cd PATH_TO_WHOLECELL_MASTER_DIR
addpath(PATH_TO_WHOLECELL_MASTER_DIR)
setWarnings()
setPath()
runSimulation('runner','koRunner','logToDisk',true,'outDir',
    SIMULATION_OUTPUT_DIR,'jobNumber',JOB_NUMBER,'koList',
    CELL_ARRAY_OF_GENE_CODES_TO_KO)
```

Figure 5: A Matlab script used to run a gene knock-out simulations using the simulation runner sub-class created in Supplementary Figure 4.

```
#!/bin/bash

# COMMENTS

## Job name
#PBS -N JOB_NAME

## Resource request
#PBS -l nodes=NO_OF_NODES:ppn=NO_OF_CORES,walltime=HH:MM:SS
#PBS -q QUEUE_NAME

## Job array request
#PBS -t MIN_ARRAY_NUMBER-MAX_ARRAY_NUMBER

## designate output and error files
#PBS -e /PATH/TO/DIR/TO/SAVE/STDERR/FILE
#PBS -o /PATH/TO/DIR/TO/SAVE/STDOUT/FILE

# print some details about the job
echo "The␣Array␣ID␣is:␣${PBS_ARRAYID}"
echo Running on host ‘hostname‘
echo Time is ‘date‘
echo Directory is ‘pwd‘
echo PBS job ID is ${PBS_JOBID}
echo This job runs on the following nodes:
echo ‘cat $PBS_NODEFILE | uniq‘

# load required modules
# e.g. module load apps/matlab-r2013a
echo "Modules␣loaded:"
module list

# code to be excuted for each array job should go here
```

Figure 6: A template submission script for a cluster with a PBS/Torque queueing system like BC3.

```
#!/bin/bash -login

# COMMENTS

## Job name
#SBATCH --job-name=NAME_OF_JOB

## What account the simulations are registered to
#SBATCH -A ACCOUNT_NAME

## Resource request
#SBATCH --ntasks=NUMBER_OF_TASKS
#SBATCH --time=D-HH:MM:SS
#SBATCH -p QUEUE_NAME

## Job array request
#SBATCH --array=MIN_ARRAY_NUMBER-MAX_ARRAY_NUMBER

## designate output and error files
#SBATCH --output=/PATH/AND/NAME/TO/SAVE/STDOUT.OUT
#SBATCH --error=/PATH/AND/NAME/TO/SAVE/STDERR.ERR

# print some details about the job
echo "The Array TASK ID is: ${SLURM_ARRAY_TASK_ID}"
echo "The Array JOB ID is: ${SLURM_ARRAY_JOB_ID}"
echo Running on host `hostname`
echo Time is `date`
echo Directory is `pwd`

# load required modules
# e.g. module load apps/matlab-r2013a
echo "Modules loaded:"
module list

# code to be excuted for each array job should go here
```

Figure 7: A template submission script for a cluster with a Slurm queueing system like BG.

In addition to the simulation data, another SQLite3 database was created to store biological information about the whole-cell model of *M. genitalium* - this called static.db and a copy can be found in the SI.zip file that accompanies this document. The database contains information about the genes, RNAs, proteins, protein monomers, protein complexes, and reactions within the model. This machine-readable format allows biological data to be more easily integrated into algorithms, results, and analysis.

Both ko.db and static.db have accompanying python libraries to aid in using the databases through Python - these are called ko_db.py and staticDB.py, respectively and can also be found in the SI.zip file that accompanies this manuscript.

The genetic algorithm will need to be able to perform gene knock-out simulations using the whole-cell model of *M. genitalium* - the most recent version can be downloaded from GitHub at `https://github.com/CovertLab/WholeCell` and it is assumed that this is contained somewhere on the hub in the directory *wholecell-master*. The recommended way to perform a gene knock-out experiment in the whole-cell model of *M. genitalium* is by creating a simulation runner subclass to pass the desired gene knock-out combination and a seed for the pseudo-random number generator in Matlab (Dr Jonathan Karr, personal communication November 2015) - see Supplementary Figure 4 for code used. A simulation can then be run using the code in Supplementary Figure 5.

Computer clusters normally have some kind of queueing system. PBS/Torque and Slurm are two common queuing systems which are used by BC3 and BG, respectively. A template for the submission script used is presented for BC3 (Supplementary Figure 6) and BG (Supplementary Figure 7). These may act as general templates for other PBS/Torque or Slurm clusters, respectively - alternative queuing systems will require a submission script that is a specific to that queueing system.

#### 2.2 Creating a connection class to a specific cluster using the PyGDS framework

All connection instances using the PyGDS framework must come from a class that inherits from the `Connection` class of the base_connections.py module. Since PyGDS assumes that all computers used run on Linux, the biggest difference with clusters is the queueing system used to run jobs. Two of the most common queueing systems are PBS/Torque or Slurm, and an example of each can be found in the `Bc3` and `Bg` classes from the connections.py module, respectively - the reader is encouraged to view these classes whilst reading through this section.

A parent class can be sub-classed in Python by importing the base class (i.e. `from base_connections import Connection`) and then passing the class as a parameter when defining the child class (i.e. `class Bc3(Connection):` or `class Bg(Connection):` ). The user is reminded that the base class must be initialised within the initialisation of the child class, thus requiring all the parameters required by the parent class. Additionally, the child class must define all abstract methods of the parent class.

<u>Initialisation</u>:
The child `Connection` classes have the additional initialisation variables:

- A path to a directory where all simulation output should be saved.
- A path to a directory where all files/data needed to submit jobs to the cluster should be saved.
- A path to the *WholeCell-master* directory that is necessary to run simulations using the whole-cell model of *M. genitalium*.

<u>Abstract methods:</u>
These classes do not dictate any abstract methods; however, the `Connection` class dictates that all child classes must have the `checkQueue` and `checkDiskUsage` methods.

The `checkQueue` method takes a job number as a parameter and returns all entries in the queuing system that have that specified job number. The PBS/Torque and Slurm clusters use different queuing systems and so the implementation of the `checkQueue` method are different for the PBS/Torque clusters (i.e. in the `Bc3` class) and the Slurm clusters (i.e. in the `Bg` class).

The `checkDiskUsage` method returns the user's disk usage. BC3 has a custom command, `pan_quota` , for this and so uses this command to return the percentage of disk available and used. To implement this on the BG is not straightforward because there is no equivalent to the `pan_quota` command and using multiple shared file systems makes it more complicated than it would be on a normal PC. Since this method is not used by the genetic algorithm or the job management classes we can not create the method. However, the parent class requires that the method be created and so we simply create a method of that name with the `pass` command and so it exists but does nothing when called[2].

<u>Instance methods:</u>
The `createStandardKoSubmissionScript` is a method that creates an executable PBS/Torque or Slurm submission script (the `Bc3` class or `Bg` class, respectively) that will submit a batch of gene knockout simulations using the whole-cell model of *M. genitalium*. All the method needs to know is

[2]The user should be careful when using this class on other algorithms or job management systems because they might need this function and thus cause unexpected bugs.

- The local path and file name of the submission script that it will create.
- The name of the job that will be sent to the cluster queuing system.
- The number of gene knock-out sets in this batch of jobs.
- The remote path and file name of a file that contains all the names of each of the gene knock-out sets. [3]
- The remote path and file name of a file that contains all the gene codes of each of the gene knock-out sets. [4]
- The number of times that the user wishes each gene knock-out set to be repeated.
- The path to the *WholeCell-master* directory.
- The path where the simulation data output will be stored.
- The path and file name where the simulation's standard-out should be saved.
- The path and file name where the simulation's standard-error should be saved.

The method checks that a feasible number of simulations has been given and then splits the simulations across array jobs and cores within an array job such that it gets through the cluster queue as quickly as possible.

When a job is submitted to a cluster queuing system, a job number is normally returned to standard-out. The `getJobIdFromSubStdOut` method takes the number from the standard-out and remembers it so that the job's progress can be monitored.

The file `static.db` is an SQLite3 database that acts as the central authority on data related to *M. genitalium* and its whole-cell model (see section 2.1). Children of the `Connection` class may want to query this database for various reasons, and so there are four instance methods that relate to this.

There is a Python library in the same directory as static.db that makes querying the database easier. If one wants to send a raw SQLite3 query to static.db, then the `sendSqlToStaticDb` method will do this and return the result. If one wants to use any of the other functions in the library, then the `useStaticDbFunction` can be used.

---

[3] These names must be unique and must be in the same order as the gene knock-out sets file - there is one name per line.

[4] These sets of codes must be in the same order as the gene knock-out sets names file - there is one comma-separated set of gene codes per line.

The `convertGeneCodeToId` method converts a tuple of gene codes into gene IDs.

The `getGeneInfo` method takes a tuple of gene codes and returns a dictionary containing the following attributes taken from the supplementary information of [1]:

- gene code;
- gene type (e.g. mRNA or rRNA etc);
- gene name;
- gene symbol;
- functional unit of gene product;
- deletion phenotype according to [1];
- essential in model according to [1];
- essential in experiment according to [1].

This method is not strictly necessary for the algorithm to run but is simply added for convenience to the user and as an example of how an algorithm may gain access to biological data.

#### 2.3 Constructing a job manager

The job manager has no base class to inherit from. However, it must take child classes of the `Connection` class that are able to create a submission script to run all possible simulations required by the algorithm. In this case, the algorithm will only require gene knock-out simulations in the whole-cell model of *M. genitalium* and so the `createStandardKoSubmissionScript` method of the `Bc3` and `Bg` classes satisfy this condition. Additionally, the algorithm will pass instructions of which gene knock-out simulations to run in the form of a dictionary whose keys are unique names to identify the simulation parameters and the value is the parameter values (i.e. a tuple of gene codes to knock-out).

The job manager libraries are kept in one module `batch_jobs` which contains two classes, `JobSubmission` and `ManageSubmission` . The job manager libraries do not have a rigid structure defined by abstract classes since this structure is defined in the computer communication and the algorithm libraries. Whilst there is no inheritance structure between the `JobSubmission` and `ManageSubmission` classes, it is important to note that they are intricately linked by the fact that the `ManageSubmission` class requires an instance of the `JobSubmission` class in order to be created. This structure enables the user to change the post-simulation data processing through only the `ManageSubmission` class (e.g. it becomes easy to change the data storage solution used by the system).

#### 2.4 The `JobSubmission` class

The `JobSubmission` class holds everything needed to submit a batch of jobs to a computer cluster.

Initialisation:
The class needs to be initialised with:

- A name for the submission.
- A child class that inherits from the `Connection` class of the `base_connection` with a method, known by the algorithm, that is able to create a submission script to perform all possible simulations required by the algorithm.
- A Python dictionary whose keys are unique names and the values are tuples of genes codes where each code represents a gene to knock-out.
- A base path on the remote computer where the simulation data should be saved.
- A base path on the remote computer where the simulation standard-error files should be saved.
- A base path on the remote computer where the simulation standard-out files should be saved.
- A base path on the remote computer where the files needed to run the simulations should be saved.
- The number of times the user wants each gene knock-out set to be repeated.
- A path to the *WholeCell-master* directory where the whole-cell model of *M. genitalium* is stored.

Instance methods:
The `createUniqueJobName` method creates a unique name so that files can be created and stored locally. It needs to be unique because an algorithm instance may want to submit more jobs than can be handled in one `JobSubmission` instance and so there will be multiple, very similar instances running at the same. In order to make sure that similar instances do not interfere with each other's files, a directory name that is guaranteed to be unique is needed.

Files often need to be created and transferred to the remote computer before a job can be submitted to the cluster. `prepareForSubmission` is the method that does this.

The `submitJobToCluster` method submits the job to the cluster, records the time and job number, and then deletes any temporary files created locally for the submission.

##### 2.5 The `ManageSubmission` class

The `ManageSubmission` class submits a `JobSubmission` instance to a cluster using the `submitJobToCluster` method and then monitors its progress in the queue. When the job is finished, it converts the raw simulation output into Pandas DataFrames, updates ko.db, and remembers the average growth rate and division time of all the simulations.

Initialisation:
The class needs to be initialised with:

- An instance of the `JobSubmission` class.
- Sometimes an algorithm needs to pass information specific to only that algorithm, and so there is a class variable that this can be passed to if necessary.
- The class automatically submits the job contained in the `JobSubmission` instance which can be a problem for unit testing and so a variable is passed to tell the class whether to initialise normally or in test mode.

Instance methods:
The `prepareDictForKoDbSubmission` method creates a dictionary that is designed to be recognised by the `ko_db` module and so can be used to update the `ko.db` database. This returns the dictionary that will be submitted to the database, but all data related to the simulations in the job submission will not be filled in yet - it will only contain the common data like the details of the person who submitted the job, the details about the cluster, and the time that the job was submitted.

The `prepareSimulationDictForKoDbSubmission` method goes to the directory of a specific simulation on the remote computer to open the Pandas DataFrame, extract the average growth rate and the time step when the `pinchedDiamter` variable was first zero (i.e. the time of division - if the cell did not divide then it returns the number zero). It then returns a dictionary where the key is the gene knock-out set that defines the genome of the organism, and the value is the average growth rate and division time of that organism.

The `monitorSubmission` method watches every simulation related to the job submission as it progresses through the queuing system by checking the queue after the first hour, followed by 15-minute intervals after that. Occasionally some jobs might get lost in the queuing system or the simulation crashes, and this method will account for these events. When this method finds that simulations have finished, it converts the data from state-*.mat files to Pandas DataFrames stored in .pickle files - this is done in parallel using `ProcessPoolExecutor` from the `concurrent.futures` module. As each simulation finishes and the data is converted, the average growth rate and division time are also retrieved into a dictionary using the `prepareSimulationDictForKoDbSubmission` method - these are added to a dictionary that is stored as a class variable and so once all the simulations are completed the relevant data can be found all in one place. When the whole job submission is finished, the data is converted and the growth and division time data is collected, then the method updates ko.db using the `KoDb` class of the ko_db.py module which can be found on a drive that is directly accessible from the cluster login nodes.

The `convertDataToPandas` method goes to the directory of the simulation data output and converts all the raw data from state-*.mat files into Pandas DataFrames stored in .pkl files using the Python package Pickle. It is worth noting that the .mat files are read by the `File` method of the `h5py` package. However, it was found that occasionally it threw an error whilst trying to read the file even though Matlab had no problem. The standard version of .mat file that Matlab uses is 7.0, which is a compressed version, and it appeared that the uncompressed version, 7.3, did not cause this error. In order to avoid these errors, code in the whole-cell model was modified so that it saves the files in version 7.3 rather 7.0 - lines 916 and 932 of /.../WholeCell-master/src/+edu/+stanford/+covert/+cell/+sim/+util/DiskLogger.m.

#### 2.6 Creating an algorithm using the PyGDS framework

This section will describe creating an algorithm class on the PyGDS framework using a genetic algorithm as an example - full code can be seen in the `GeneticAlgorithm` class of the multigeneration_algorithm.py module.

Supplementary Figure 8 shows how each generation of the `GeneticAlgorithm` is executed - to see how this fits in with the entire process it should be compared to Supplementary Figures 1 and 2.

<u>Initialisation</u>:
In addition to the parameters needed to initialise the `MGA` class the `GeneticAlgorithm` class requires

- The maximum number of fit individuals that are allowed to survive each generation.
- The number of times each simulation needs to be repeated.
- The number of children needed in each generation.
- A path indicting where all simulation data should be stored. This is relative to the cluster base path which is given by the connection class.
- The probability that a mutation occurs whilst creating a child.
- The name of a function (that exists in the child class) that can be called to get a dictionary that contains all the gene codes and IDs that make up the genome.

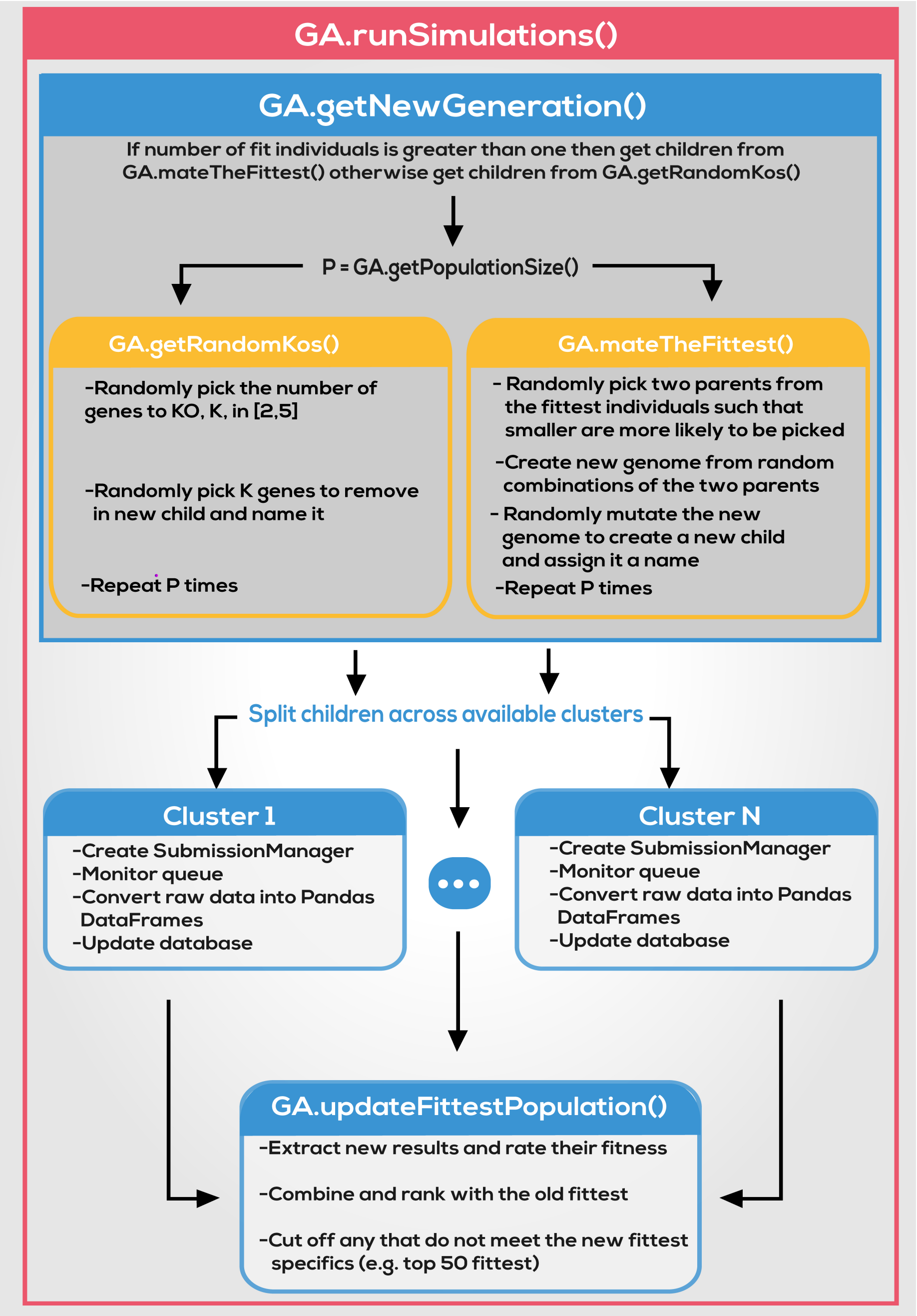


Figure 8: This diagram show how the `runSimulations` method is implemented in the `GeneticAlgorithm` class. Red boxes contain everything that happens within the `runSimulations` method. Blue boxes contain any significant methods or classes called within the `runSimulations` method. Yellow boxes contain significant methods or classes called within the blue boxes. Here GA represents the `GeneticAlgorithm` class.

Abstract methods:
The `MGA` class defines three abstract methods that need to be defined in any child classes, `getGenerationName` , `getNewGeneration` , and `runSimulations` .

The `getGenerationName` method returns the string 'genN' where 'N' is the current generation number.

The `getNewGeneration` method decides what method to call to generate the next generation of children. For the genetic algorithm, this calls the `mateTheFittest` method if the number of fit individuals is greater than one. Otherwise, it calls the `getRandomKos` method.

The `runSimulations` method defines the algorithm over one generation. For `geneticAlgorithm` this means using the `getGeneration` method to get all the child genomes for the next generation of simulations as well as the number of clusters available. It then splits the children evenly over all the clusters and creates `JobSubmission` instances for each set.

Parallel computing is required to create the `ManageSubmission` classes. Python executes all lines of code sequentially and waits for each line of code to finish before executing the next. The sequential nature of code execution means that normal code will submit the first batch of jobs to cluster-1 and then wait for the whole `ManageSubmission` process to finish before submitting the second batch of jobs to cluster-2. This sequential use of the clusters defeats the point of having multiple computing facilities, and so a parallel solution was created by using the `multiprocessing` library to `map` each job to their respect clusters. Due to the amount of time it takes to convert the simulation data output to Pandas DataFrames and the fact that it is not uncommon for lots of simulations to finish at a similar time, the `ManageSubmission` class executes the `convertDataToPandas` method in parallel as well. It is now clear that the process running the `ManageSubmission` class is already a child process from the parallelised mapping in the `runSimulations` method. Unfortunately, the `multiprocessing` library does not allow child processes to spawn new child processes, and so the more popular library was dropped and replaced by the `futures` module of the `concurrent` library.

Once all the simulations for this generation have completed then the `runSimulations` method passes all the finished `ManageSubmission` instances to the `updateFittestPopulation` method in order to *learn* what happened in the current generation.

Instance methods:
The `getPopulationSize` method returns the desired population of children for this generation.

The `getRandomKos` method finds out what genes can be knocked-out out from the genome from class variables and uses the `getPopultionSize` method to find out how many children need to be created. It then uses these to create the desired number of children, each with a random number of genes knocked out in the range $[2, 5]$. The number of genes knocked out and which genes are knocked out are both picked from a uniform distribution. Each child name is made up of two parts, the first part is 'ko' and the second part is a number that starts equal to 1 and increments by one every time a new child is created so that each child has a unique name in the generation. When a new generation starts, the name counter goes back to 1. This method returns a dictionary where the keys are the names of the children, and the values are the gene codes of the genes that need to be knocked out - this dictionary will be referred to as the *child name to gene knock-out set dictionary.*

The `mateTheFittest` method creates all the children for the next generation. The children are created by mimicking natural selection and sexual reproduction by randomly selecting two parents from the fittest individuals found so far (natural selection) and creating a child by mixing the genomes of the two parents (sexual reproduction). It starts by making a copy of all the fittest individuals. The genome of individual $i$, $\Theta_i$, can be represented by the set of all genes knocked-out of the wild-type, $K_i$. The fittest individuals, $F$, are a set of knock-out that represent all the individuals that have survived up until that point, $F = \{K_i\}$. Each of the fittest individuals is assigned a fitness score which is the number of knock-outs, $L = \{l_i \mid (l_i = |K_i|) \wedge (K_i \in F)\}$, and thus can define the probability of picking individual $K_i$ as

$$\mathbb{P}(X = K_i) := \frac{l_i}{\sum_{l_j \in L} [l_j]}. \quad (1)$$

Two individuals, $K_i$ & $K_j$, are randomly picked from the fittest set to mate using equation 1. These are converted into the genome representation and will become the parents of a new child, $P^1 = \Omega(K_i) = \{\theta_1^1, \theta_2^1, \ldots, \theta_{|\Gamma|}^1\}$ and $P^2 = \Omega(K_j) = \{\theta_1^2, \theta_2^2, ..., \theta_{|\Gamma|}^2\}$[5]. In order to create the child a random number in the range $x \in [1, |\Gamma|]$ is uniformly picked, this indicates how much of the child genome will be made up of parent-1. A subset of size $x$ is uniformly picked from $P^1$, $C^{P^1} \subset P^1$, and the remaining genes will come from parent-2, $C^{P^2} \subset P^2$. $C^{P^1}$ and $C^{P^2}$ are combined to make a new genome, $C = C^{P^1} \cup C^{P^2} = \{\theta_1^{k_1}, \theta_2^{k_2}, \ldots, \theta_{|\Gamma|}^{k_{|\Gamma|}}\}$ where $k_m = 1$ if taken from parent-1 or $k_m = 2$ if taken from parent-2. A mutation probability is passed at initialisation of the algorithm, and this used to determine which children receive a random mutation from a uniform distribution. The number of genes to be mutated is picked randomly from a custom[6] method based on an exponential distribution

[5]The user defines the total gene set, $\Gamma$, and thus its size when creating an instance of the algorithm. The gene set may be the entire wild-type genome or some subset of that which excludes genes that the user does not want to knock-out. For this case study, $|\Gamma| = 358$, which is the number of the characterised protein-coding genes minus one that tends to cause the simulations to crash.

[6]It was desired that the number of genes to knock-out be variable in order to enable large

```
number_of_gene_mutations = 0
        while number_of_gene_mutations == 0:
                number_of_gene_mutations = int(np.around(np.random.
                    exponential(2)))
```

Figure 9: Code segment showing how the modified exponential distribution is calculated. The while-loop means that a 0 value will never be created, the `np.around` method performs standard rounding on the result, and the `int` method converts the data type from float to integer ( `int` truncates all decimal places rather than rounding them and so rounding them first results in fewer loops).

with parameter two (see code segment 9 for the code). 10,000 samples were taken from the modified exponential distribution, and a histogram of the data can be seen in Supplementary Figure 10. The sample minimum, maximum, mean, and standard deviation are 1, 20, 2.54050, and 2.00049, respectively. Once the number of genes is known then that number of genes are selected randomly from a uniform distribution and then *flipped* by adding genes that were knocked-out or knocking-out genes that were present (i.e. $\theta_i$ is a binary variable and so can be *flipped* by adding 1 modulo 2, $\theta_i \to (\theta_i + 1) mod 2$). The process of creating children is then repeated until enough children are created for the new generation.

The `updateFittestPopulation` method takes the simulation results from a completed `SubmissionManager` instance and extracts all individuals that produced a dividing cell. The dividing cells are then combined with the current fittest individuals and ranked so that the smallest genomes are at the top and the largest at the bottom. The algorithm class is initialised with a maximum number of fit individuals, $M$, and so the top $M$ individuals are taken from the new list and set as the new fittest individuals.

<u>Instance methods for all child classes:</u>
These are methods deemed generally useful and automatically get put into all child classes of the `MGA` class and will always be identical in implementation. These will not be defined again in the other child classes.

The `random_combination` method takes a Python iterable (e.g. a list) and the desired size and then picks a random subset of that size from a uniform

and small random mutations to occur. However, the larger the number of mutations the larger the chance of killing the cell and so a uniform distribution with a large range will result in mostly killing the cell, but a small range means that larger mutations can *never* be tested. A desirable distribution would be a negative exponential distribution - this enables both large and small numbers of mutations. However, small numbers of mutations are more likely with the probability of larger numbers of mutations exponentially decaying. The problem with the negative exponential distribution is that it is continuous in values starting from 0 whereas we desire discrete integers starting from 1.

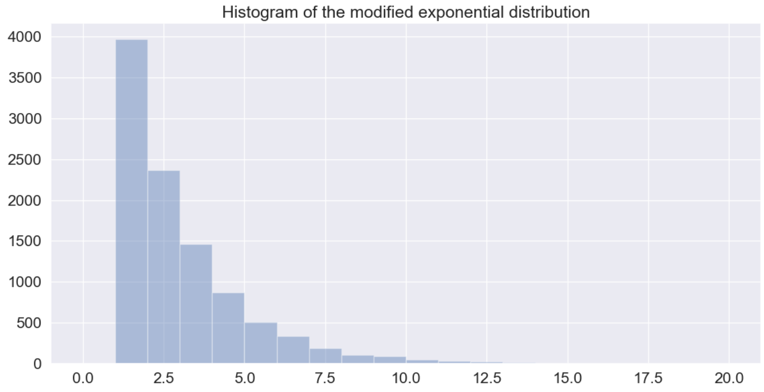


Figure 10: Histogram of the modified exponential distribution. 10,000 data points were sampled from the modified exponential distribution to create this histogram. The data had a minimum value of 1, a maximum value of 20, a mean value of 2.54050, and a standard deviatation of 2.00049.

distribution which is then returned to the user.

The `random_pick` method is the same as the `random_combination` method except it picks the iterable elements from a distribution defined by an iterable of probabilities passed by the user.

The `getJr358Genes` method returns a tuple of gene codes. These gene codes are defined as all of the protein-coding genes that are characterised in the whole-cell model of *M. genitalium* minus one that tends to crash the simulation.

The `getDictOfJr358Codes` method returns a dictionary where the keys are gene codes returned by the `getJr358Genes` method and the values correspond to the ID used in our databases. This method takes an instance of the `Connection` class and uses that connection to get IDs directly from `static.db` so that all users are working off the same data source.

The `invertDictionary` method takes a dictionary and, assuming that the keys and values share a bijective relationship, returns a dictionary where the keys and values are swapped.

The `createIdxToIdDict` method takes a dictionary that converts gene codes into gene IDs and converts that into a dictionary that converts genome indexes into gene IDs.

The `convertIdxToGeneId` method takes a list of genome indexes and returns a corresponding list of gene IDs

The `convertGeneIdToCode` method takes a list of gene IDs and returns a corresponding list of gene codes.

#### 2.7 Running the genetic algorithm

Having set up the /.ssh/config file as specified in section 1.1; created a sub-class of the `Connection` class to enable the communication between the users hub and cluster(s); set up the databases and corresponding communication modules on the cluster(s); and made all the PyGDS modules available to an instance of Python3+ on the hub, it will be possible to run a genetic algorithm to reduce the genome of the *M. genitalium* whole-cell model on the user's cluster(s).

Supplementary Figure 11 shows a Python script that could be used to run the genetic algorithm described in this document. In a Python3 virtual-environment, a user can start the genetic algorithm with `python genetic_algorithm_runfile_example.py` . Due to the long running time of these massive *in-silico* experiments, it is advised that a user runs the Python script in a terminal multiplexer (e.g. GNU screen or tmux) so that it is possible to log out of a remote connection with the hub and then log back in and carry on the session. In addition to this, it is useful to be able to both see the progress of the algorithm in real-time whilst also being able to look at the standard-out and standard-error in a text file. In order to fulfil both of these requirements a user can start the algorithm with the following command `python genetic_algorithm_runfile_example.py 2>&1 | tee path/to/outfile.out` .

### 3 Other modifications

This case study and Rees and Chalkley et al.[2] have presented examples of how to use PyGDS to run two different genome reduction experiments on the whole-cell model of *M. genitalium* on two different high-performance computer clusters. It is possible to modify this to include different different data management requirements, models and design objectives. Section 2.3 described creating the job management classes for data management specific to the genome design group at the University of Bristol; however it should be relatively easy to recreate that data management system or modify it for other systems. Adapting this code to use different models and design objectives would not be as straight forward as say running existing algorithms on different clusters or adding a new algorithm. This is because the code changes need to be made throughout all the different modules. To run an algorithm on a different model changes would need to be made to the child connection classes and the job management classes so that they know how to submit simulations using the new model to the cluster. Varying different parameters in the model (e.g. not gene knock-outs) would require making the same changes as well as changes to the `MGA` sub-class since

```
# load classes to be used
from multigeneration_algorithm import GeneticAlgorithm
from connections import Bc3 # this example uses the Bc3 class but
    the user needs to load a connection sub-class that will work on
     their cluster

# details about the user
cluster_user_name = 'USER_NAME'
ssh_config_alias = 'SSH_ALIAS'
path_to_key = 'PATH_TO_SHH_KEY'
name1 = 'USER_FIRST_NAME'
name2 = 'USER_SURNAME'
email = 'USER_EMAIL_ADDRESS'
# create instances of the connection classes that the user wishes
    to run simulations on
bc3_connection = Bc3(cluster_user_name, ssh_config_alias,
    path_to_key, name1, name2, email)
dict_of_cluster_instances = {'bc3': bc3_connection} # This
    dictionary will be passed to the algorithm class and can
    contain multiple cluster instances
dict_for_checkStop = {'max_generation': [100]} # This will tell the
     algorithm to stop when it gets to 100 generations
MGA_name = 'GA_full_run_with_seed_2019_05_15' # This must be a
    unique name for the in-silico experiment (if not unqie then the
     simulations will throw an error a stop)
max_no_of_fit_individuals = 100 # This is the maximum number of fit
     individuals allowed to mate to create the next generatio
reps_of_unique_ko = 1 # This is the number of times each simulation
     needs to be repeated

# create an instance of the GeneticAlgorithm class
genetic_algorithm = GeneticAlgorithm(dict_of_cluster_instances,
    dict_for_checkStop, MGA_name, max_no_of_fit_individuals,
    reps_of_unique_ko, generation_num_to_gen_size_dict = {0: 600,
    -1: 200}) # generation_num_to_gen_size_dict tells it to create
    600 children for generation 0 and 200 children for all other
    generations
genetic_algorithm.run() # start the genetic algorithm
```

Figure 11: A example of a Python script that will use a genetic algorithm to reduce the genome of the *M. genitalium* whole-cell model - actual file can be found in genetic_algorithm_runfile_example.py from the SI.zip file that accompanies this document.

this needs to know what parameters use.

### References


[1] Jonathan R Karr, Jayodita C Sanghvi, Derek N Macklin, Miriam V Gutschow, Jared M Jacobs, Benjamin Bolival, Nacyra Assad-Garcia, John I Glass, and Markus W Covert. A whole-cell computational model predicts phenotype from genotype. *Cell*, 150(2):389–401, jul 2012.

[2] Joshua Rees, Oliver Chalkley, Sophie Landon, Oliver Purcell, Lucia Marucci, and Claire Grierson. Designing Minimal Genomes Using Whole-Cell Models. *bioRxiv*, page 344564, mar 2019.

[3] Darrell Whitley. A genetic algorithm tutorial. *Statistics and Computing*, 4(2):65–85, jun 1994.

[4] Thomas Back. *Evolutionary algorithms in theory and practice : evolution strategies, evolutionary programming, genetic algorithms.* Oxford University Press, 1996.

[5] ACRC. *BlueCrystal Phase 3.* https://www.acrc.bris.ac.uk/acrc/phase3.htm.

[6] BrisSynBio. *BlueGem.* https://www.bristol.ac.uk/brissynbio/equipment/hpc/bluegem/.
